## Supplementary Information for "Melon: metagenomic long-read-based taxonomic identification and quantification using marker genes"

### Supplementary Tables

#### Table S1: Computational time and peak memory usage for wastewater samples

| | | no pre-filter | | PlusPF-8 | | PlusPF-16 | | PlusPF[b] | |
|---|---|---|---|---|---|---|---|---|---|
| influent 7.816 Gb | genome copy | 1,809 | | 1,801 | | 1,800 | | 1,797 | |
| | species richness | 2,101 | | 2,098 | | 2,099 | | 2,097 | |
| | number of filtered reads | - | | 8,312 | | 10,212 | | 14,988 | |
| | mean genome size (Mb) | 4.322 | | 4.304 | | 4.300 | | 4.292 | |
| | ARG abundance (copies per cell)[a] | 0.551 | | 0.553 | | 0.553 | | 0.554 | |
| | real time (sec)[c] | 2,056 | 967 | 2,271 | 1,027 | 2,279 | 1,151 | - | 1,272 |
| | peak resident set size (GB)[c] | 10.629 | 17.860 | 10.665 | 18.143 | 17.294 | 17.986 | - | 77.694 |
| effluent 5.158 Gb | genome copy | 1,348 | | 1,336 | | 1,331 | | 1,315 | |
| | species richness | 1,704 | | 1,697 | | 1,700 | | 1,696 | |
| | number of filtered reads | - | | 16,602 | | 29,300 | | 54,774 | |
| | mean genome size (Mb) | 3.826 | | 3.789 | | 3.757 | | 3.715 | |
| | ARG abundance (copies per cell)[a] | 0.507 | | 0.511 | | 0.512 | | 0.519 | |
| | real time (sec)[c] | 1,341 | 671 | 1,496 | 885 | 1,495 | 867 | - | 929 |
| | peak resident set size (GB)[c] | 10.893 | 13.649 | 10.902 | 13.227 | 17.067 | 17.454 | - | 77.325 |

[a] Excluding multidrug ARGs.
[b] Not tested with MacBook Pro due to insufficient memory.
[c] Measured with GNU 'time'. Red: MacBook Pro 2021, with Apple M1 Max, 64 GB memory, and macOS Sonoma 14.0. Black: Lab-scale workstation, with 2 × Intel Xeon Silver 4210R CPU 2.40GHz (10 cores, 20 threads), 512 GB memory, and Ubuntu 20.04 LTS.

**Table S2: Performance of PHMMs at different scales of threshold scores**

| | scale | precision | recall | $F_{0.5}$-score | $F_1$-score | RPGF[a] |
|---|---|---|---|---|---|---|
| bacteria | 0.50 | 0.969 | 0.931 | 0.954 | 0.942 | 42 |
| | **0.75** | **0.983** | **0.961** | **0.977** | **0.969** | **45** |
| | 1.00 | 0.984 | 0.958 | **0.977** | 0.968 | **45** |
| archaea | 0.50 | 0.920 | 0.677 | 0.800 | 0.734 | 21 |
| | **0.75** | **0.936** | **0.703** | **0.809** | **0.749** | **23** |
| | 1.00 | 0.906 | 0.639 | 0.750 | 0.688 | 19 |

[a] Number of RPGFs with PHMM's $F_{0.5}$-scores greater than 0.99.

#### Table S3: Statistics of mock and wastewater samples

| platform | mock | sample | read | Gb[a] | length | | | | quality score | |
|---|---|---|---|---|---|---|---|---|---|---|
| | | | | | longest | shortest | mean | median | mean | median |
| ONT | D6300 | S1 | 2,503,848 | 16.619 | 185,010 | 1,000 | 6,637 | 3,947 | 10.899 | 11.037 |
| | | S2 | 2,119,258 | 9.514 | 51,733 | 1,000 | 4,489 | 3,673 | 12.085 | 12.044 |
| | | S3 | 336,330 | 3.486 | 211,938 | 1,000 | 10,365 | 3,879 | 13.187 | 13.320 |
| | D6331 | G1 | 5,650,723 | 27.891 | 50,325 | 1,000 | 4,935 | 4,569 | 13.742 | 14.243 |
| | | G2 | 1,570,234 | 6.458 | 51,516 | 1,000 | 4,112 | 3,581 | 16.987 | 17.321 |
| | | G3 | 713,445 | 3.587 | 38,817 | 1,000 | 5,028 | 4,171 | 19.045 | 19.365 |
| | - | influent[b] | 1,091,772 | 7.816 | 374,946 | 1,000 | 7,158 | 6,295 | 18.320 | 18.677 |
| | | effluent[b] | 1,118,502 | 5.158 | 91,897 | 1,000 | 4,611 | 3,443 | 18.211 | 18.539 |
| PacBio | MSA-1003 | - | 2,418,889 | 20.544 | 21,547 | 1,001 | 8,493 | 8,310 | 38.768 | 34.784 |
| | D6331 | - | 1,978,476 | 17.993 | 39,601 | 1,003 | 9,094 | 8,078 | 45.521 | 39.623 |

[a] Gigabase pair ($10^9$ base pairs).
[b] Two spike-ins species *Allobacillus halotolerans* and *Imtechella halotolerans* were removed before calculation.

**Table S4: Error rates of models trained on mock samples**

| mock | sample | mismatch | insertion | deletion | **total** |
|---|---|---|---|---|---|
| D6300 | S1 | 0.040 | 0.015 | 0.025 | **0.080** |
| | S2 | 0.026 | 0.024 | 0.032 | **0.082** |
| | S3 | 0.021 | 0.014 | 0.017 | **0.052** |
| D6331 | G1 | 0.017 | 0.011 | 0.019 | **0.046** |
| | G2 | 0.010 | 0.006 | 0.013 | **0.029** |
| | G3 | 0.009 | 0.006 | 0.007 | **0.022** |

### Supplementary Figures

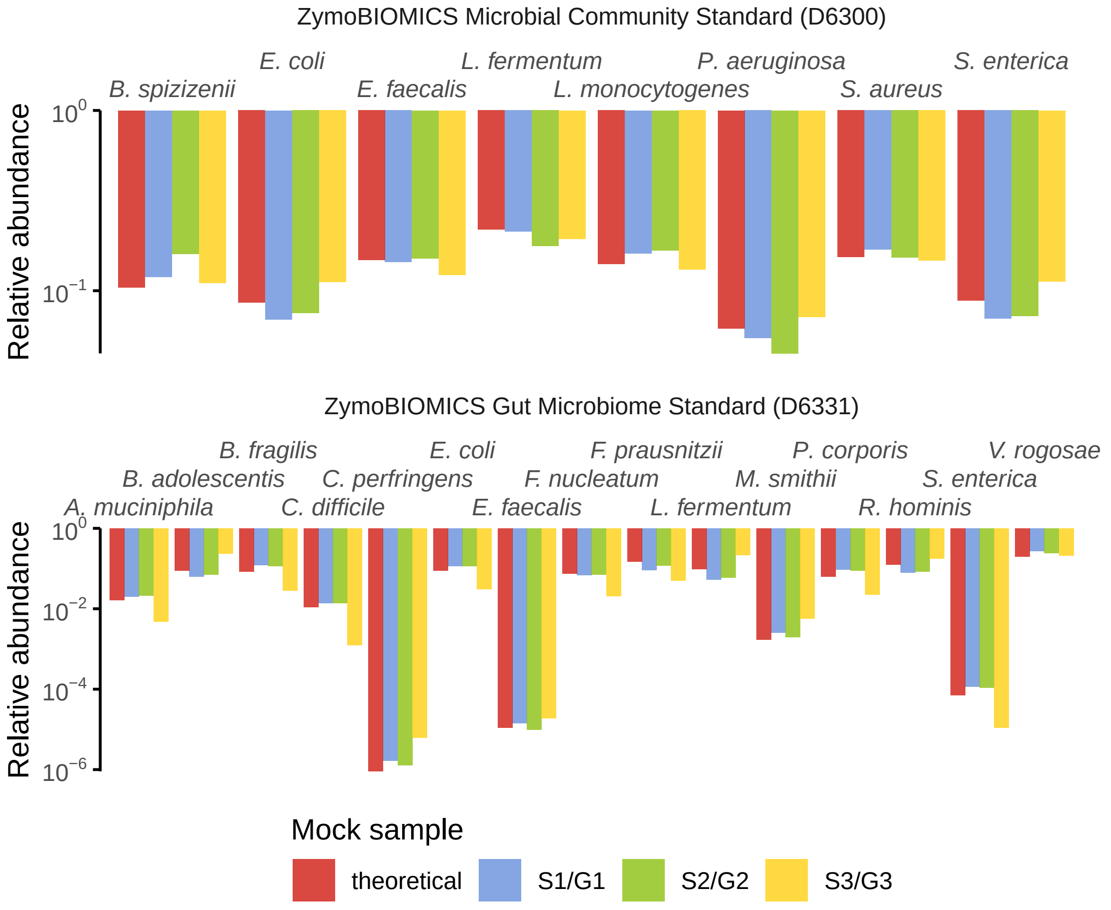

**Figure S1: Comparison between theoretical and expected taxonomic composition of mock communities D6300 and D6331.** Theoretical relative abundances (in terms of genome copies) were provided by ZymoBIOMICS. Expected relative abundances were obtained by mapping reads to their associated reference genomes. Yeasts were excluded in both theoretical and expected abundances. Plasmids were not counted while computing expected relative abundances. *Bacillus subtilis* and *Lactobacillus fermentum* have been renamed to *Bacillus spizizenii* and *Limosilactobacillus fermentum*, respectively.

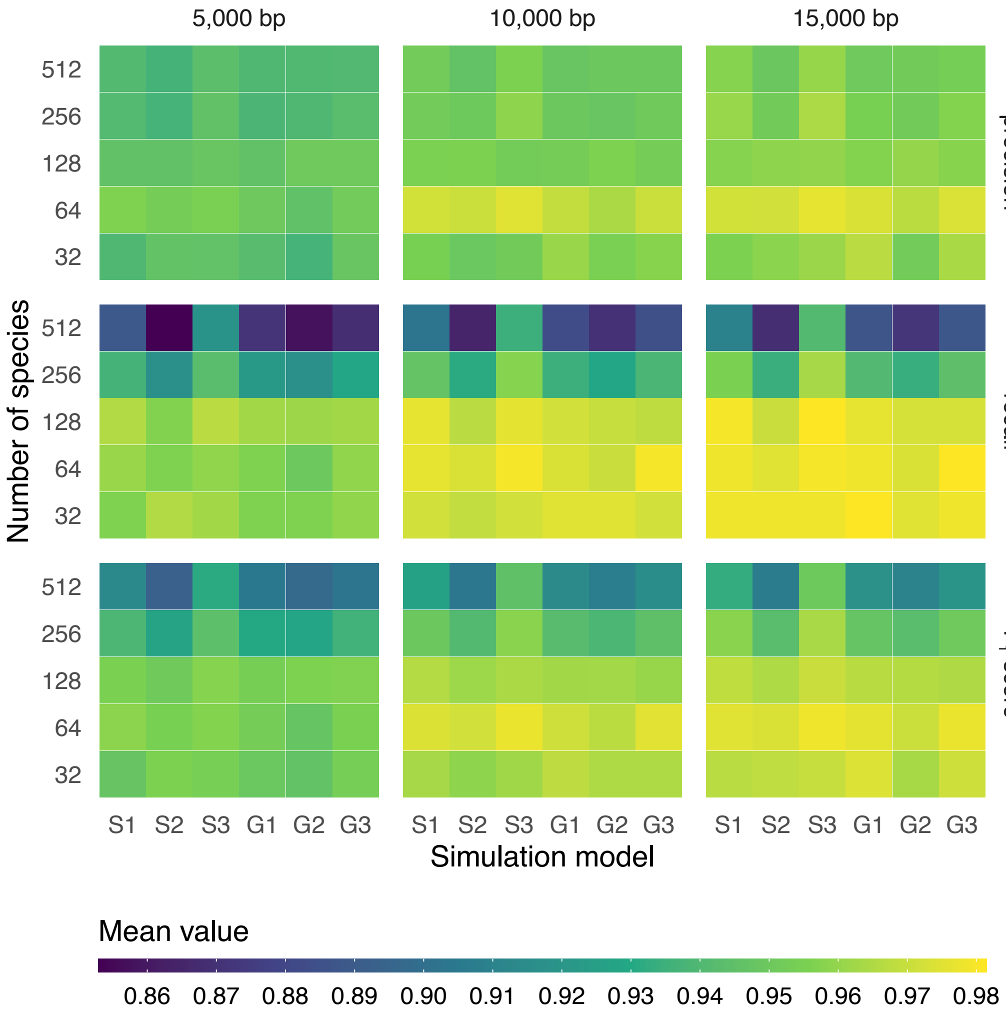

**Figure S2: Performance of Melon at different length cutoffs of flanking regions.** Models used for simulation were trained using six mock samples (S1–3 and G1–3). Each combination of simulation models and numbers of species contains ten randomly generated profiles. Colors represent the mean values of these profiles.

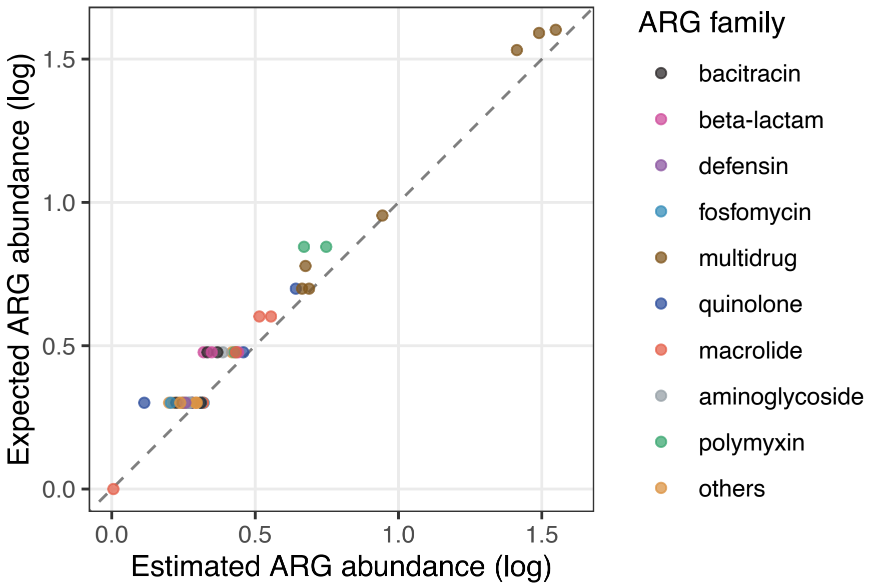

**Figure S3: Estimated species-level ARG abundances of mock sample S3.** Estimated species-level ARG abundances (expressed as "copies per cell", assuming one genome copy per cell) were computed by normalising the estimated copies of ARGs by the estimated genome copies provided by Melon. Expected ARG abundances were obtained from the reference genomes given by ZymoBIOMICS. Colors indicate the families of mostly observed ARGs. ARG abundances are shifted by "+1" before $\log_{10}$-transformation to avoid zero entries.

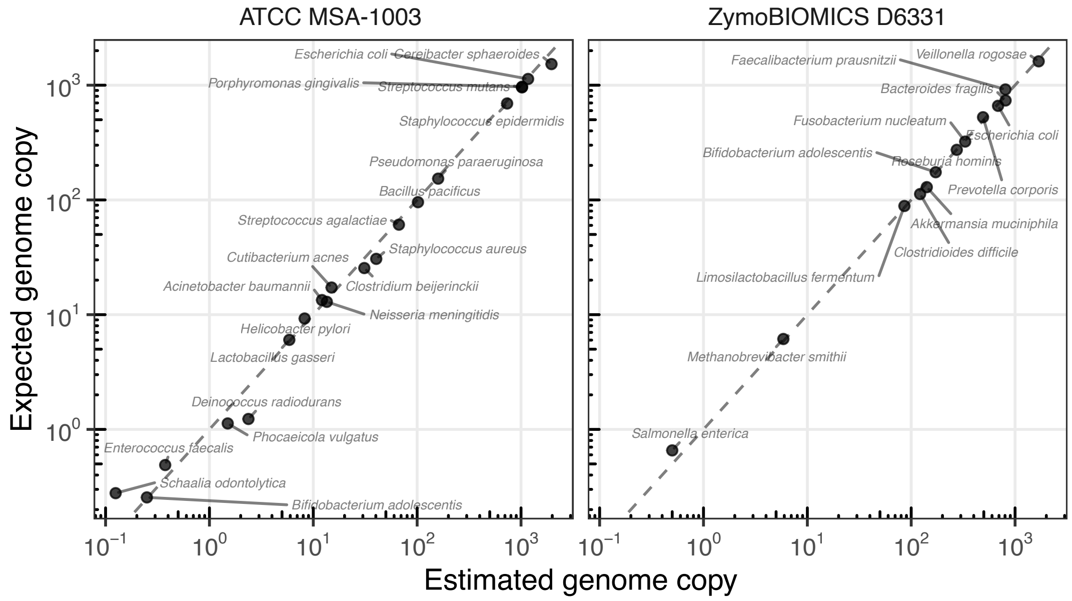

**Figure S4: Estimated genome copies of PacBio samples.** Estimated genome copies were returned by Melon. Expected genome copies were obtained by mapping PacBio reads to their respective reference genomes. Reads that mapped to yeasts or remained unmapped were filtered out. Species are labelled with texts. *Pseudomonas aeruginosa* has been renamed to *Pseudomonas paraeruginosa* and *Lactobacillus fermentum* to *Limosilactobacillus fermentum.*
